# Improved sensitivity and resolution of ATAC-seq differential DNA accessibility analysis

**DOI:** 10.1101/2022.03.16.484118

**Authors:** Ahmed Ali Sheikh, Alexandre Blais

**Affiliations:** University of Ottawa, Faculty of Medicine, Department of Biochemistry, Microbiology and Immunology, 451 Smyth Road, Ottawa, ON, K1H 8M5, Canada; Éric Poulin Centre for Neuromuscular Disease, Ottawa, ON, Canada; Ottawa Institute of Systems Biology, Ottawa, ON, Canada; University of Ottawa Centre for Inflammation, Immunity and Infection (CI3), Ottawa, ON, Canada

## Abstract

Eukaryotic genomes are packaged into chromatin, and the extent of its compaction must be modulated to allow several biological processes such as gene transcription. The regulatory elements of expressed genes are typically in relatively accessible chromatin, and several studies have revealed a reliable correlation between the abundance of mRNA transcripts and the degree of DNA accessibility at the regulatory elements of their coding genes. In consequence, the genome-wide profiling of DNA accessibility by methods such as ATAC-seq can help in the study of gene regulatory networks by serving as a proxy for gene expression and by helping identify important gene cis-regulatory elements and the trans-acting factors that bind them. The predominant approach used to identify differentially accessible genomic loci from ATAC-seq data obtained in two conditions of interest is comparable to that employed in RNA-seq gene expression profiling studies: accessible regions are identified through peak calling and treated like “genes”, then sequenced DNA fragments (originating from two neighboring transposase insertion events) that overlap them are counted and subjected to abundance modeling, which then allows to identify those that have a significant difference between the two conditions. We reasoned that this approach could be improved in terms of sensitivity and resolution by introducing two changes: bypassing peak calling, using instead a genome-wide sliding window quantification approach, and counting transposase insertion sites, instead of fragments originating from two neighboring insertion sites. We present the development of this approach, which we term “widaR”, for Window- and Insertion-based Differential Accessibility in R, using a murine skeletal myoblast differentiation dataset. Reproducible R code is provided.

## Introduction

The accessibility of DNA within chromatin is an important feature of genome organization in eukaryotes: gene expression levels are significantly correlated with DNA “openness” because the transcriptional machinery typically requires unhindered access to its template. This is illustrated among others by several transcription factors (TFs) recruiting chromatin remodeling enzymes as co-activators, by pioneer factors providing DNA access to other gene regulatory factors through direct or indirect disruption of nucleosomes, and by gene regulatory elements being the sites of incorporation of replication-independent histone variants facilitating access to DNA (*1–4*). Like mRNA expression profiles, DNA accessibility patterns provide information about cell identity (*5–7*). Additionally, genome-wide DNA accessibility profiles also carry useful information on the potential activity of trans-acting regulatory elements (*5*), which constitutes an advantage over transcriptomic profiles when regulatory mechanisms are the focus of attention.

Several experimental methods exist to identify sites of high DNA accessibility: DNase I hypersensitive sites sequencing (DNase-seq (*5*)), formaldehyde-assisted isolation of regulatory elements coupled to sequencing (FAIRE-seq (*8*)), and assay of transposase-accessible chromatin using sequencing (ATAC-seq (*9*)). Due to its ease and amenability to low or ultra-low input cell amounts, ATAC-seq has become the leading method at time of writing (Figure S1). It uses a hyperactive Tn5 transposase (Tnp) that will cut DNA and insert custom-designed adapters at sites where chromatin in sufficiently accessible. DNA flanked by two insertion sites is amplified by PCR and sequenced at high throughput. A commonly used method to identify genomic loci of differential accessibility (DA sites) between two or more conditions consists in identifying regions of substantial accessibility in at least one condition and counting high-throughput sequencing reads (or read pairs) aligning to these regions; those with significantly different counts between conditions are considered DA sites (*10, 11*).

However, certain assumptions made for RNA-seq differential mRNA abundance testing are not necessarily valid for ATAC-seq analysis. Others have previously reported the strong impact of the normalization scheme used (*12*). An example is whether it is appropriate to normalize on the count matrix of peak-overlapping reads (*i.e*. how many reads fall in peaks, in each library). In modern RNA-seq datasets, most reads can be assumed to come from legitimate mRNA species that the experimenter wishes to quantify. Instead, in ATAC-seq and ChIP-seq, the fraction of reads in peaks (FRiP) may represent a minority of all reads: the ENCODE projects indicates a minimum acceptable value of 30% for the FRiP score for ATAC-seq datasets (*13*). In this context, normalizing by depth estimation using the peak-overlapping reads count matrix would disregard most reads. This may be a problem with certain datasets where the samples have uneven FRiP scores. Alternatives proposed for ChIP-seq and/or ATAC-seq are to normalize on the entire library, on the fraction of the reads that are not in peaks (*i.e*. background reads), or to use methods such as loess that can deal with trended bias (*12, 14–18*).

Likewise, assumptions valid for the analysis of ChIP-seq for punctate binding TFs may turn out invalid for the analysis of ATAC-seq data. ChIP-seq is performed with an antibody against a single antigen, such as a TF or histone covalent modification mark. Immunoprecipitated chromatin is sequenced and one can assume that the site of TF binding lies, on average, in the middle of the sequenced molecule of DNA (*19*). The result is that the true site of TF binding is generally at the summit of the peaks (the location of maximum coverage signal) and it is generally expected that if differential binding exists between two sample types, the difference will have a certain homogeneity across the region called as peak.

ATAC-seq breaks these assumptions because chromatin accessibility to Tnp insertion is the consequence of potentially multiple biological factors (such as presence or absence of nucleosomes, transcription machinery components, or TFs) that cannot be unmixed straightforwardly. The result is broad peaks that may have differential accessibility in only a subregion of the peak (Figure 1). Thus, an issue with two-step DA site identification approaches that start with peak calling is that the resolution of DA site identification will be limited by the genomic length of the starting peaks. With peaks that are relatively broad and encompassing multiple local summits of signal strength, narrower local regions of DA may be obscured by the general trend of accessibility at the level of the entire peaks. The same phenomenon and challenge were noted before in the case of ChIP-seq experiments for histone modification marks, which tend to be fairly broad but have differential occupancy only in subregions of peaks (*20*).

**Figure 1.**
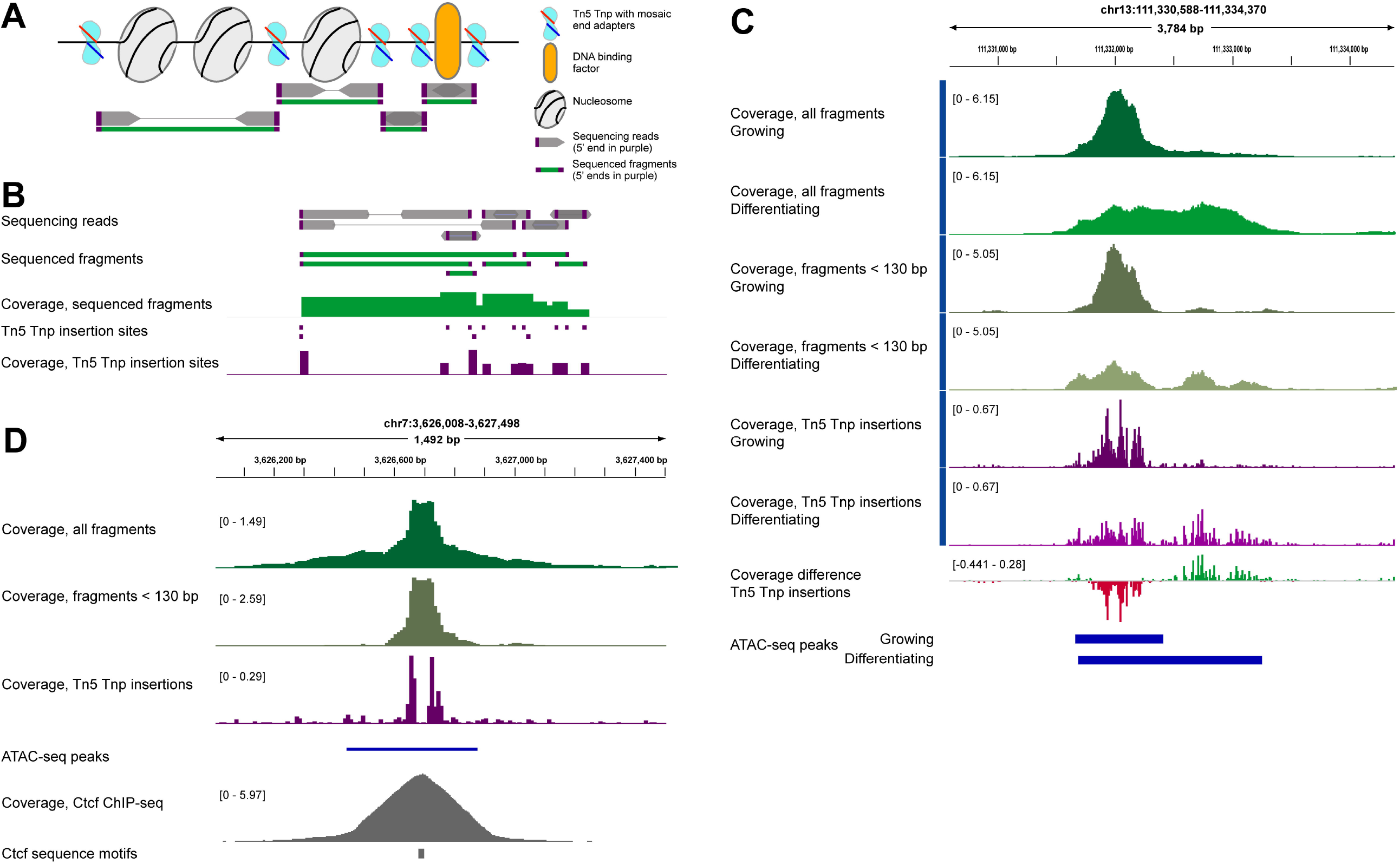
Relationship between Tn5 Tnp insertion sites and sequencing fragments, and impact on ATAC-seq data interpretation. **A)** Schematic representation of transposase insertion into chromatin and resulting sequencing reads (gray), fragments defined by read pairs (green) and transposase insertion sites (purple), illustrating how the read 5’ ends more faithfully represent the true location of where Tn5 Tnp gained accessibility to DNA Thin gray lines represent non-sequenced DNA sequence intervening between some read pairs. Not drawn at scale. **B)** Example of ATAC-seq sequencing data from a BAM file visualized as read pairs. The data genomic coverage, in 10 base pair bins, for fragments and for insertion sites are shown as histograms of matching colors. **C)** Example of a genomic locus with significant ATAC-seq signal giving rise to more than one summit and where the difference in DNA accessibility is not homogeneous. Comparing the signal in growing and differentiating myoblasts, the peaks identified overlap but are not equal. Further, the ATAC-seq signal for differentiating cells decreases in the left-hand side part of the region but increases on the right-hand side part. This is reflected by coverage difference between the two samples that is highly variable across this region, going from negative (red) to positive (green) difference values. **D)** Example of a genomic locus where considering Tn5 Tnp insertion sites provides higher resolution and uncovers biologically meaningful information unseen when Tn5 Tnp fragments are examined. At this location, binding of the Ctcf protein renders DNA inaccessible to Tn5 Tnp, creating a typical “TF footprint”.

In some studies, any genomic locus unoccupied by nucleosomes, *i.e*. nucleosome-free, has been considered “accessible DNA”. This has been achieved by restricting analyzed sequencing fragments (in BAM files) to the read pairs that match a certain maximum fragment length, for example sub-nucleosomal fragments (*9*). While perfectly suitable to address certain biological questions, this approach has the disadvantage of reducing the amount of available information in a dataset (by disregarding all read pairs above that insert size cut-off) and makes debatable assumptions about the true level of accessibility of DNA in between the two Tnp insertion sites. For example, two Tnp insertion sites that are at 100 base pairs from each other cannot be separated by a nucleosome, yet they may still be separated by a DNA-binding protein that alters DNA accessibility (Figure 1); such has been shown to be the case for CTCF and other transcription factors (*15*). We reason that in some contexts, it may be best for DA analysis to consider Tnp insertions without biasing the analysis towards nucleosome presence or absence. This could include the study of pioneer transcription factors, which are generally thought to bind compact (*i.e*., nucleosomal) DNA and lead to the opening of chromatin, or when studying proteins that are not amenable to transcription factor footprinting (*e.g*., general DNA binding proteins like those of the RNA pol II machinery without DNA binding sequence preference).

We reasoned that a one-stage (*i.e*., peak calling-independent) approach with higher resolution would help circumvent these problems. Here, we compare the performance of two main approaches to analyze ATAC-seq: the common two-stages method starting with peak calling, and a one-stage method based on sliding window quantification of signal. Each method was employed to quantify Tn5 Tnp insertion sites or fragments. Further, the sliding window method was compared with THOR, another peak calling-independent method based on hidden Markov models (HMM) and previously shown superior to other one- and two-stages approaches in ChIP-seq applications (*20*). To test these methods, we employed a well-defined biological system for which abundant genomic data exist for validation purposes: the model of murine C2C12 skeletal myoblast differentiation into myotubes (*21*). These cells are immortal and easy to cultivate, yet they recapitulate the key steps of myogenesis, including irreversible cell cycle arrest, cellular fusion, terminal differentiation, and activation of several transcription factors implicated in making muscle (*22–25*). In the present study, chromatin accessibility in C2C12 cells was compared in two states: in proliferation conditions or 24 hours following the induction of differentiation. The result of extensive testing shows that the method we developed, which we named *widaR*, is the best performing one.

## Results

To analyze ATAC-seq data with sliding windows, we used the R/Bioconductor package *csaw* for a few reasons. First, it is well-documented and highly flexible, allowing end-users to adapt the workflow to the specific nature of their experiment. In particular, it natively allows analysis of either fragments (defined by two neighboring Tn5 Tnp insertions and reflected in the sequencing results as properly paired sequencing mates) or Tn5 Tnp insertion sites (reflected in the data as the 5’ end of every read, independent of pairing with mates). Further, csaw is seamlessly integrated with the edgeR analysis framework, a widely-adopted and well-documented Bioconductor package for differential expression analysis which provides options for differential signal modeling and allows the use of strategies to remove unwanted batch effects within samples (*26*).

The myoblast dataset was obtained using the OMNI-ATAC method on four biological replicates of myoblasts in growth phase (referred to as “Grow”, in figures) and three biological replicates of cells after 24 hours in differentiation medium (called “Diff” in figures). At this stage, the myoblasts have exited the cell cycle and initiated the expression of muscle differentiation and function genes like Myogenin, various myosin and troponin isoforms (*27–29*) and this is known to coincide with chromatin remodeling and changes in DNA accessibility (*30–32*). The OMNI-ATAC method was selected for its ability to reduce mitochondrial DNA reads. On average, approximately 5% of all uniquely aligning reads mapped to chrM (Table S1).

In all comparisons, the ATAC-seq data were analyzed to discover DA regions between cell states (more or less accessible in growing or differentiating myoblasts). Key settings for the methods tested are summarized in Table 1. We initially performed the analysis in three different ways: **I)** csaw with Tnp insertion sites; **II)** csaw with sequenced fragments; **III)** MACS2 peak calling followed by featureCounts summarization of sequenced fragments and edgeR differential analysis. Methods **I** and **II** employed windows of 50 base pairs that are immediately adjacent and non-overlapping (*i.e*., distance of 50 base pairs center-to-center), as this value was deemed a reasonable compromise to increase resolution and sensitivity compared to peaks without draining computational resources. Window size and distance parameters are easily adjusted, and the impact of changing their values is discussed below. We note that method **II** is equivalent to approach V tested by Reske et al. (*12*), except that a window width of 300 bp was used in their study.

**Table 1.** Summary of methods tested.

| Name | Genomic regions | Counted elements | DA testing method | c.f. Figure # |
| --- | --- | --- | --- | --- |
| Method I | Sliding windows 50 bp | Read 5' ends | csaw/edgeR | Figure 3, Figure 4 |
| Method Ib | Sliding windows 25 bp | Read 5' ends | csaw/edgeR | Figure 6 |
| Method Ic | Sliding windows 100 bp | Read 5' ends | csaw/edgeR | Figure 6 |
| Method II | Sliding windows 50 bp | Seq. fragments (all) | csaw/edgeR | Figure 4 |
| Method IIb | Sliding windows 50 bp | Seq. fragments (length < 130 bp) | csaw/edgeR | Figure 4 |
| Method III | Fragment peaks | Seq. fragments (all) | featureCounts/edgeR | Figure 3 |
| Method IIIb | Fragment peaks | Seq. fragments (length < 130 bp) | featureCounts/edgeR | Figure 4, Figure S8 |
| Method IIIc | Fragment peaks | Read 5' ends | featureCounts/edgeR | Figure 6 |
| Method IIId | Insertion peaks | Seq. fragments (all) | featureCounts/edgeR | Figure S8 |
| Method IIIe | Insertion peaks | Seq. fragments (length < 130 bp) | featureCounts/edgeR | Figure S8 |
| Method IV | Sliding windows 50 bp | Read 5' ends | THOR (HMM) | Figure 6 |

Method **III** relies on quantification of fragments in ATAC-seq peaks called using MACS2 on sequencing fragments because this is the method we found most often used in the literature. Information from biological replicates was combined to arrive at sets of consensus peaks for ‘Grow’ and ‘Diff’ samples, and a union of both consensus sets was generated and used in DA testing (Figure S2).

First, we compared methods **I, II** and **III** before and after TMM normalization using global analysis with RLE (relative log expression (*33*)) and PCA (principal component analysis) plots. This yielded comparable results between methods and showed that the samples cluster primarily by cell state, regardless of method (Figure 2, right). The fragment-based methods (**II** and **III**) tended to show higher variance explained in PC1 compared to insertion-based (method **I**). TMM normalization, with counts from windows with signal 5 times above the genome-wide average (methods **I** and **II**) or within-peaks counts (method **III**), efficiently reduced sample heterogeneity and brought the medians very close to zero (Figure S2, left). Using LOESS normalization instead of TMM did not yield an appreciable difference (Figure S3). Various parameters for removal of unwanted variation were also tested; the results of these empirical tests led us to select RUVs with a k value of 2, for each method (Figure S4).

**Figure 2.**
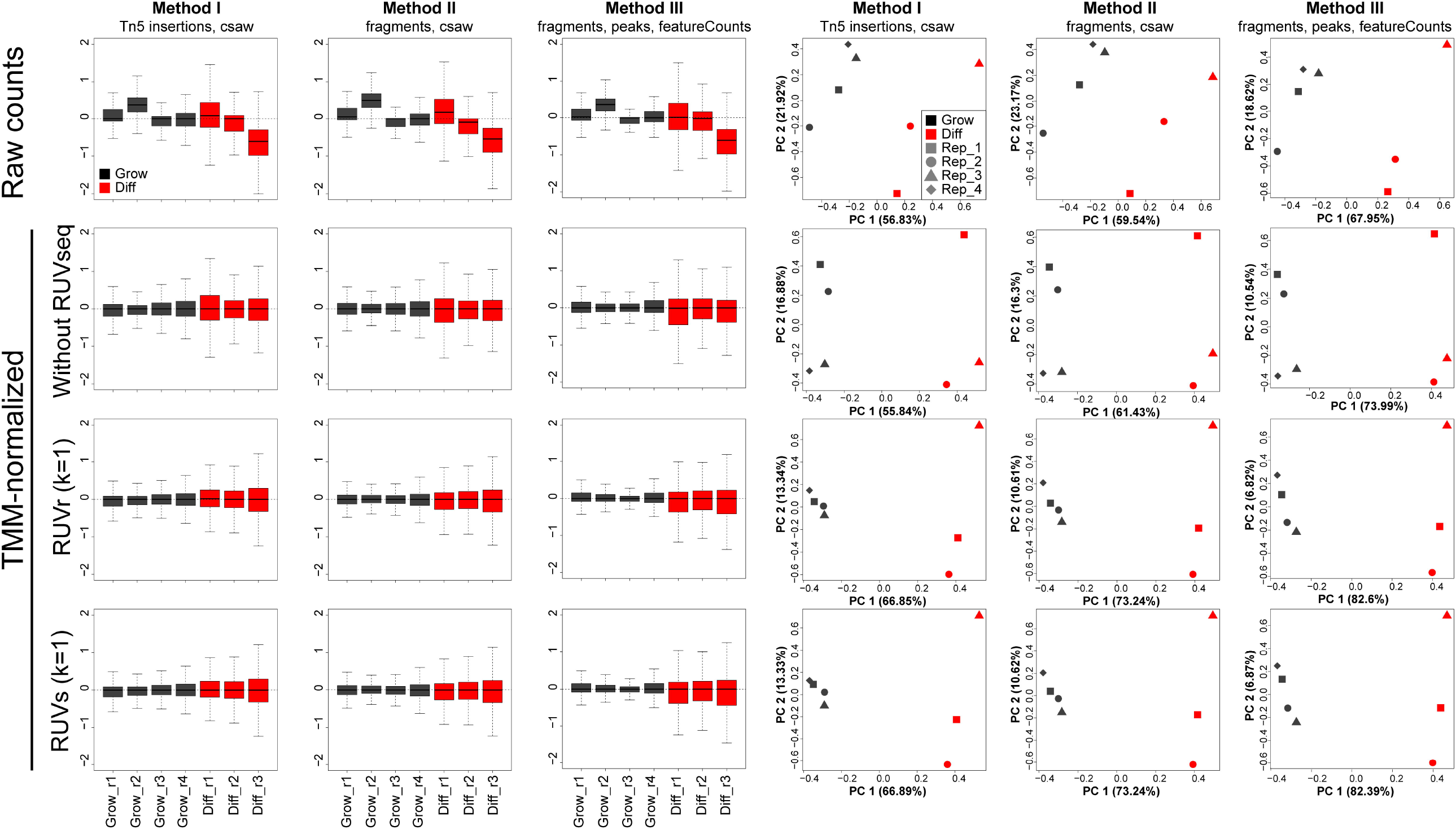
Comparison of signal quantified by the different methods. RLE and PCA plots are shown with the three methods on raw data, TMM-normalized data, and after applying RUVseq batch effect removal with the RUVr or RUVs algorithms.

### Comparing the three base methods

Windows or peaks with DA were identified, using cut-offs of Benjamini-Hochberg-corrected p values lower than 0.05 and log_2_ fold-change values above 1 or below −1. Because certain DA genomic regions may comprise multiple significant windows (especially since windows are smaller than peaks (Figure S2), such windows were merged if they were immediately adjacent (Figure S5). The results of the comparison are shown in Figure 3, Figure S6, Figure S7 and Table S2. While method **I** identified 37,891 significant DA regions, method **II** identified 35,517 and method **III** 18,866 (Figure 3A). As expected, the DA regions identified by the three methods overlap to a significant extent, but they also identified unique regions: method **I** identified the most, with 10,852 unique DA regions, while method **III** identified the fewest (Figure 3B).

**Figure 3.**
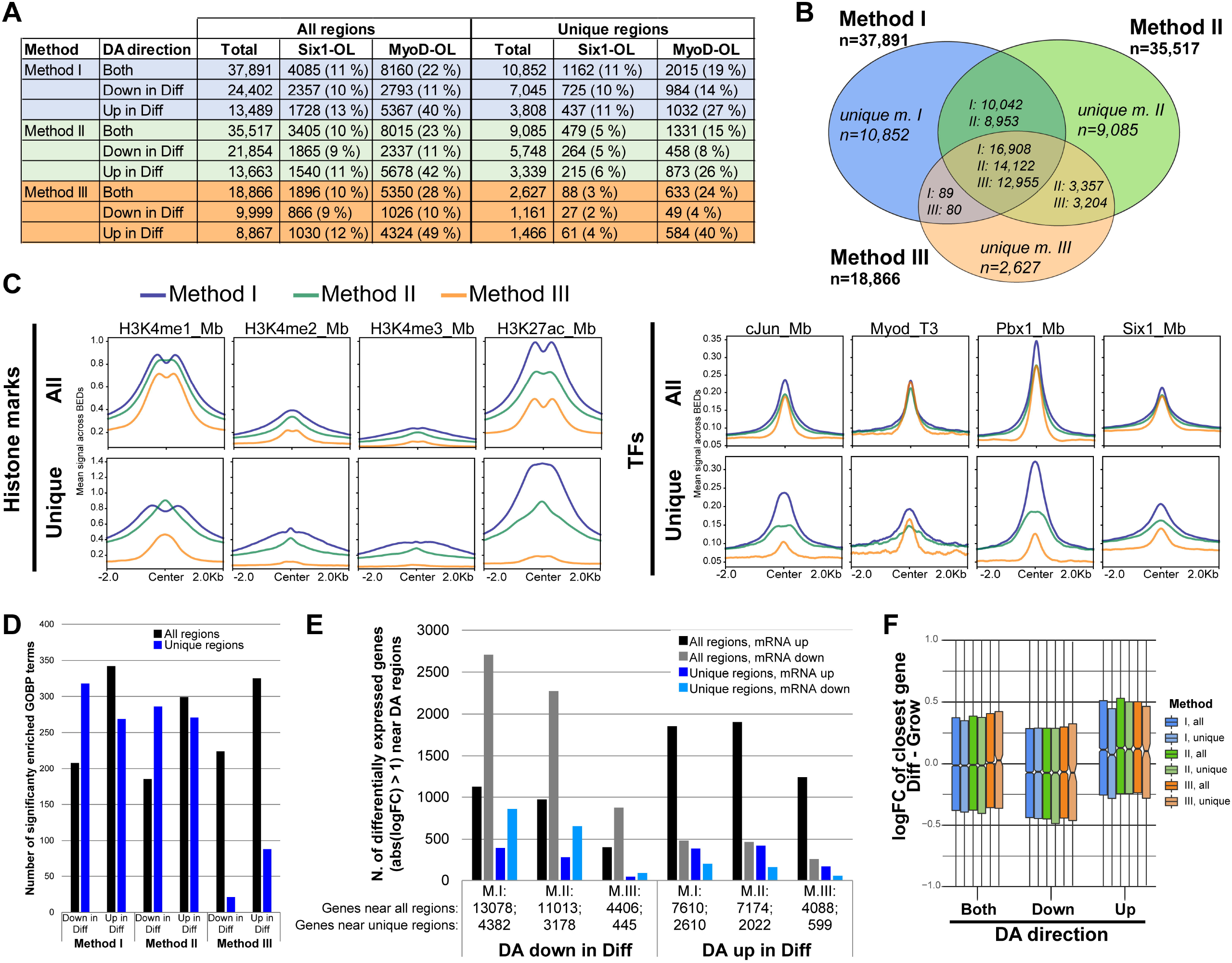
Comparison of DA site identification results obtained with methods I, II and III. In this comparison, counting Tnp insertions in sliding windows (method **I**) was compared with counting all sequenced fragments in sliding windows using csaw (method **II**) or in peaks using featureCounts and edgeR (method **III**). **A)** Table summarizing the numbers of DA sites discovered by each method, broken down in regions that decrease or increase in accessibility during myoblast differentiation. Unique regions refer to DA regions discovered using one method but not the other two. The overlap between these regions and ChIP-seq peaks for MyoD or Six1 is also provided. With sliding window-based methods (i.e. **I** and **II**), the regions are those obtained after the merging of significant neighboring windows (*c.f*., Figure S5). **B)** Venn diagram representing the degree of overlap between the DA regions discovered by each method. Some regions found with one method may overlap two narrower regions discovered by another method. For this reason, the overlapping areas of the diagram give numbers for each method. Note that areas are not exactly proportional. **C)** Mean signal for histone marks and transcription factors for the entire sets of DA regions discovered by each method (“All”) or those unique to each method (“Unique”). Refer to Figure S7 for the actual heatmaps of genomic coverage signal. **D)** Number of GO biological process terms that are enriched (with observed over expected ratio of 2.5 or higher, and adjusted p-value of 1E-4 or lower) among the genes located near DA regions discovered by each method. Black bars represent the results for all DA regions found by a given method, and blue bars those for the regions found only by a given method. The analysis was split between DA regions with increased and with decreased accessibility. **E)** Histogram indicating the number of differentially expressed genes near DA regions identified by each method. Genes were assigned to regions as the closest gene withing 50 kb. Regions are broken down into those with lower or with higher DNA accessibility in differentiated cells. Genes are counted if their expression changes by at least two-fold in either direction. The total number of genes assigned to a group of DA regions is indicated at the bottom of the columns. **F)** Gene expression logFC value distribution, for the genes closest to DA regions.

To determine if these regions are biologically meaningful and relevant, we examined their overlap with genomic features of known importance, namely binding sites of muscle TFs and abundance of histone post-translational modification marks. The DA regions identified by method **I** overlapped the largest number of MyoD and Six1 binding sites (Figure 3A); this is most evident for the regions uniquely found by method **I**. The ChIP-seq signal for histone marks H3K4me1, H3K4me2, H3K4me3 and H3K27ac were found to be most abundant at DA regions identified by method **I** (Figure 3C); this was especially pronounced for H3K4me1 and H3K27ac, marks associated with enhancers and active regulatory elements, respectively. The ChIP-seq signal for MyoD, Six1 and two additional TFs important for muscle, c-Jun and Pbx1, was also examined, and likewise found to be most enriched at DA sites identified by method **I**.

To further establish the biological relevance of the DA regions discovered, we employed gene ontology (GO) term enrichment analysis. We reasoned that if DA regions are biologically meaningful, they are likely to represent regulatory elements of genes involved in common pathways and that this commonality will be evidenced by significant enrichment of the GO biological process terms associated to them. First, we looked at the GO term annotations of the genes closest to the DA regions and calculated the enrichment level and statistical significance for each term in each DA region set. As expected, we observed enrichment of several GO terms in the groups of regions with increased or decreased DNA accessibility (Figure 3D, “All regions” column). Interestingly, the two sliding window-based methods (**I** and **II**) performed better than the peak-based method **III** with this metric; this is particularly evident when the GO term enrichment analysis was performed with the method-specific DA regions: method **I** returned the most enriched GO terms and method **III** the fewest.

As a third evaluation of biological significance, we examined the expression of the genes located near DA regions, reasoning that DA regions should be associated to differentially expressed (DE) genes (*5, 34*). We examined the number of DA regions located near DE genes (Figure 3E). Method **I** performed comparably to method **II** and markedly better than method **III** according to this metric, discovering more DA regions associated to gene expression regulation; this is seen both with all DA regions or only those unique to a given method (Figure 3E, black and blue bars, respectively). Finally, the log_2_ fold-change value distribution of these genes was examined, which revealed that methods **I, II** and **III** perform similarly according to this metric (Figure 3F).

We conclude from these analyses that method **I** has improved sensitivity (it identifies a larger number of DA regions) and specificity (the DA regions are more enriched in biologically relevant features) compared to method **III**, the typical method employed to identify DA sites in the genome.

### Comparing the effect of restricting sequencing fragment lengths

Next, we wondered if method **II** and **III** would reach the performance of method **I** if only sequenced fragments matching sub-nucleosomal lengths were included in the analysis. This is logical because sequencing fragments less than 147 bp cannot be generated from Tnp insertion events on each side of a nucleosome, so the presence of these short fragments is indicative of nucleosome-free regions where possibly most of DA events occur. For methods **IIb** and **IIIb**, we limited the analysis (by sliding window and by counting at peaks, respectively) to fragments less than 130 bp to ensure that only transposition events from nucleosome-free regions are counted. A summary of the results is given in Figure 4. First, method **I** yielded a significantly larger number of DA regions, compared to methods **IIb** or **IIIb** (Figure 4A). Method **IIIb** (counting sub-nucleosome length fragments at peaks) is the method finding the lowest number of DA regions. This was paralleled by a larger number of regions uniquely identified by method **I** (Figure 4B). The biological relevance of the discovered DA regions with method **I** is manifested also by the superior numbers of MyoD and Six1 binding sites overlapping them (Figure 4A). Histone and TF ChIP-seq signal coverage appears equal or slightly higher with method **IIb** compared to method **I**. Again, with this criterion method **IIIb** is the weakest performer. At the gene expression level, method **I** is superior to method **II**, as it discovers a larger number of DA regions near DE genes (Figure 4E). The global trend in expression difference of genes near DA regions is comparable with all three methods, except for the fact that the few regions uniquely found by method **IIIb** are likely dominated by spurious hits with inconsistent expression difference (Figure 4F). Overall, we conclude that method **I** is superior to **IIb** and **IIIb**. Further, method **IIb** appears superior to method **II** in terms of biological relevance, but this is at the expense of sensitivity (compare Figure 4A with Figure 3A), which probably comes from the fact that a lower amount of information (read pairs filtered by fragment size) enters the analysis, reducing statistical power: restricting the analysis to only the sequencing fragments of 130 bp or less removes 60% of the read pairs on average (Table S1).

**Figure 4.**
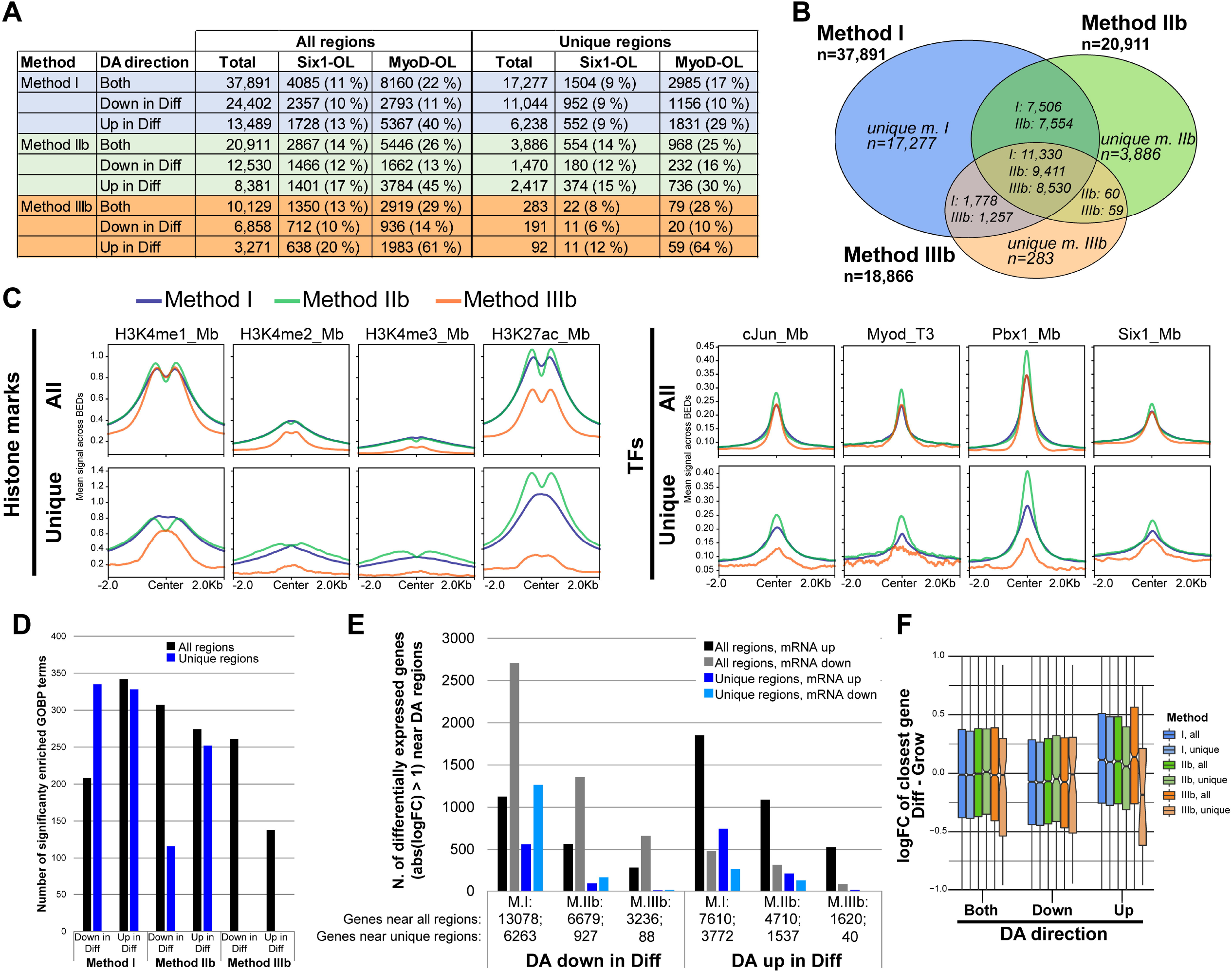
Comparison of DA site identification results obtained with methods I, IIb and IIIb. In this comparison, only sequenced fragments of sub-nucleosomal length were counted, either in sliding windows using csaw (method **IIb**) or in peaks using featureCounts and edgeR (method **IIIb**). **A)** Summary of number of all and unique DA regions discovered and overlap with MyoD or Six1 binding sites. **B)** Venn diagram showing the overlap between DA regions identified by each method. **C)** Mean signal for histone marks and transcription factor ChIP-seq. **D)** Number of GO biological process terms that are enriched among the genes located near DA regions. **E)** Number of DE genes near DA regions. **F)** General gene expression (logFC) trend for genes near DA regions.

### Comparison of Tnp insertion sites quantification methods

Finally, we compared the performance of three methods based on quantification of transposase insertions. Method **I** was compared with method **IIIc**, where Tnp insertions are quantified over peaks, and method **IV**, where THOR is used. THOR is a differential peak identification algorithm based on hidden Markov models (HMM) developed for ChIP-seq datasets (*20*). THOR is comparable to csaw in that it is a sliding window-based approach that does not require prior identification of peaks (*18*). It was developed for single-end ChIP-seq data, but we applied it to the ATAC-seq data, specifying to extend reads by 1 base pair, which had the result of only considering the Tnp insertion site on each mate independently (as with method **I**). However THOR internally applies a read strand correlation filtration step to ChIP-seq differential peak candidates (*19*), which is irrelevant for ATAC-seq data but was impossible to omit. To make it comparable to method **I**, a corrected p-value of 0.05 and log fold-change cut-off of 1 were applied to the THOR output. THOR by design combines adjacent significant windows into DA peaks, so post-hoc merging of significant regions is unnecessary. The results of these comparisons are given in Figure 5. The first notable observation is that method **IV** identified the largest number of sites with significant DA (about 10% increase over what is discovered with method **I**). Interestingly, the proportion of DA sites with decrease in ‘Diff’ over ‘Grow’ is higher with method **IV** compared to methods **I** or **IIIc** (73% compared to 64% and 53%, respectively). The reason for this is not immediately clear but could be related to the entirely different computational strategy to make DA calls (generalized linear model compared to HMM).

**Figure 5.**
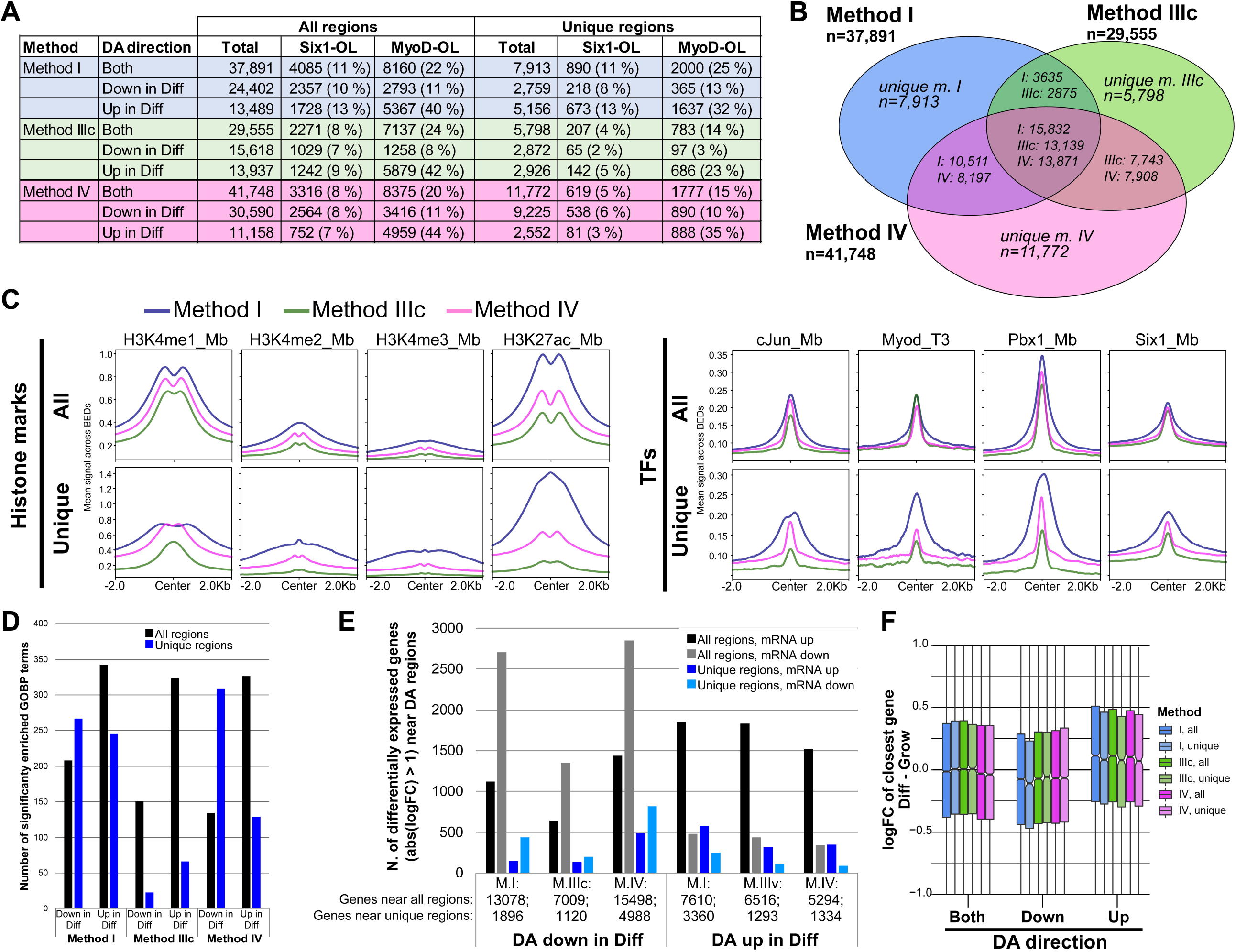
Comparison of DA site identification results obtained with methods I, IIIc and IV. This analysis compares the performance of the three methods based on Tnp insertion sites: quantified on sliding windows with csaw (method **I**), quantified over peaks using featureCounts and edgeR (method **IIIc**), or analyzed with by HMM method with THOR (method **IV**). **A)** Summary of number of all and unique DA regions discovered and overlap with MyoD or Six1 binding sites. **B)** Venn diagram showing the overlap between DA regions identified by each method. **C)** Mean signal for histone marks and transcription factor ChIP-seq. **D)** Number of GO biological process terms that are enriched among the genes located near DA regions. **E)** Number of DE genes near DA regions. **F)** General gene expression (logFC) trend for genes near DA regions.

The histone marks examined show higher signal enrichment at the DA regions identified with method **I**, and this is true both for all sites and for regions unique to each method; the superiority of method **I** is most evident with H3K27ac (Figure 5C). It was noticed that the shape of the histone mark mean signal curves is different, between methods: in particular, the H3K27ac signal curve lost its bimodality at sites uniquely discovered by method **I**. The reason for this is uncertain, but one possibility is that method **I** identifies regions that are smaller compared to the other two (Figure S9), so that they have a higher likelihood of being off the center of regulatory elements such as enhancers and thereby not being symmetrically flanked on each side by covalent mark-bearing nucleosomes. At the level of TF binding sites, the three methods appear to perform equally well (Figure 5C), though when only the regions unique to each method are examined, method **I** is clearly superior for the four TFs analyzed.

Method **IIIc** turned out to represent an improvement over method **IIIb** in terms of number of DA regions discovered and absolute numbers that overlap with Six1 or MyoD binding sites. However, the increased sensitivity seems to occur at the cost of specificity. The percentage of DA regions that overlap Six1 or MyoD binding sites decreased (Figure 5A), and the degree of coincidence with histone marks indicative of gene regulatory elements decreased: while method **IIIb** yields H3K4me1 almost identical to method **I** (Figure 4C), the mean curve for H3K4me1 with method **IIIc** is well below that of method **I** (Figure 5C).

Looking at GO term enrichment, we found that methods **I** and **IV** are comparable, and superior to method **IIIc**. Again, the DA regions uniquely found with method **IIIc** reveal the worst performance, supporting the notion that they comprise a larger number of spurious hits (Figure 5D). At the level of DE of genes near DA regions, methods **IV** appeared to perform slightly better than method **I** when all DA regions were analyzed, and both performed better than method **IIIc** (Figure 5E). However, method **I** showed a slight increase in performance of method **IV** for the DA regions that increase in accessibility during differentiation. As with the previous comparisons, no marked difference between methods was observed in the log2 fold-change values of the genes near DA regions (Figure 5F). Overall, the results of this comparison show that method **I** is slightly less sensitive compared to method **IV** but finds DA regions that tend to be more biologically relevant.

### Impact of peak calling method

We note that method **III** (and variants **IIIb** and **IIIc**) relies on calling peaks, which is traditionally performed using the entire length of sequenced fragments. However, since we noted above that accessible DNA is logically more accurately represented by Tnp insertion sites (Figure 1A), we also used MACS2 to call peaks on segments centered on the 5’ ends of both reads. Comparing peaks called using fragments or Tnp insertions, substantial differences were noted in both the number of peaks discovered and their lengths: peaks called on Tnp insertion sites are more numerous and tend to be narrower (Figure S2A to F). Further, their overlap was also examined. The two methods generally agree on what genomic regions constitute peaks of accessible DNA (Figure S2G to I). However, it was found that while the peaks called with fragments cover a large number of base pairs that are not covered by insertion peaks (14421 kb, Figure S2G), these base pairs are for the majority found in peaks that are shared with insertion peaks (Figure S2J): 96% of base pairs unique to fragment peaks are within peaks that overlap with insertion peaks and only 4% are within peaks unique to the fragments method. Instead, of the base pairs that are unique to insertion peaks, 41% are within peaks that overlap with fragment peaks and 59% are in unique peaks. These observations suggest it is probable that calling ATAC-seq peaks using Tnp insertion sites is more sensitive and affords higher resolution in identifying regions of DNA accessibility. For this reason, we additionally performed a comparison of method **III** where peaks are called from fragments (methods **IIIb** and **IIIc**) or from Tnp insertion sites (methods **IIId** and **IIIe**), and where all fragments are counted (**III** and **IIId**) or only subnucleosomal length ones (**IIIb** and **IIIe**). The results confirmed our intuition, showing that peaks based on insertions are more biologically relevant and help uncover a larger number of DA regions (Figure S8). Thus, if a peak-based DA region identification method is desired, method **IIId** would seem the best performer.

### Comparison of sliding window length and distance

Finally, we explored the effect of changing the length and center-to-center distance of the sliding windows. We compared method **I** (windows of 50 bp) with methods **Ib** (windows of 25 bp) and **Ic** (windows of 100 bp). The results are presented in Figure 6. Decreasing window length marginally increased the number of DA regions identified while increasing window length instead more severely reduced the number of DA regions (Figure 6A). With its shorter windows, method **Ib** permitted the identification of a larger number of DA regions overlapping Six1 or MyoD binding sites, and this was especially obvious with the regions uniquely found by each method (Figure 6A). At the level of histone marks and TF binding signal enrichment, these three methods performed comparably, though method **Ib** was clearly superior for H3K27ac (Figure 6C). However, at the level of the regions uniquely found by these three related methods, method **Ib** was superior for all biological features examined (Figure 6C, bottom row). This was especially noticeable for H3K27ac, where method **Ib** yielded the highest mean signal and was the only one to display the characteristic bimodal pattern. In general, using 25 bp windows performed as well or better than 50 bp windows, and this identified unique regions that are biologically relevant not found when using windows of 50 or 100 bp. We conclude that quantifying Tnp insertions in sliding windows of 25 bp identifies DA regions with the best sensitivity and specificity.

**Figure 6.**
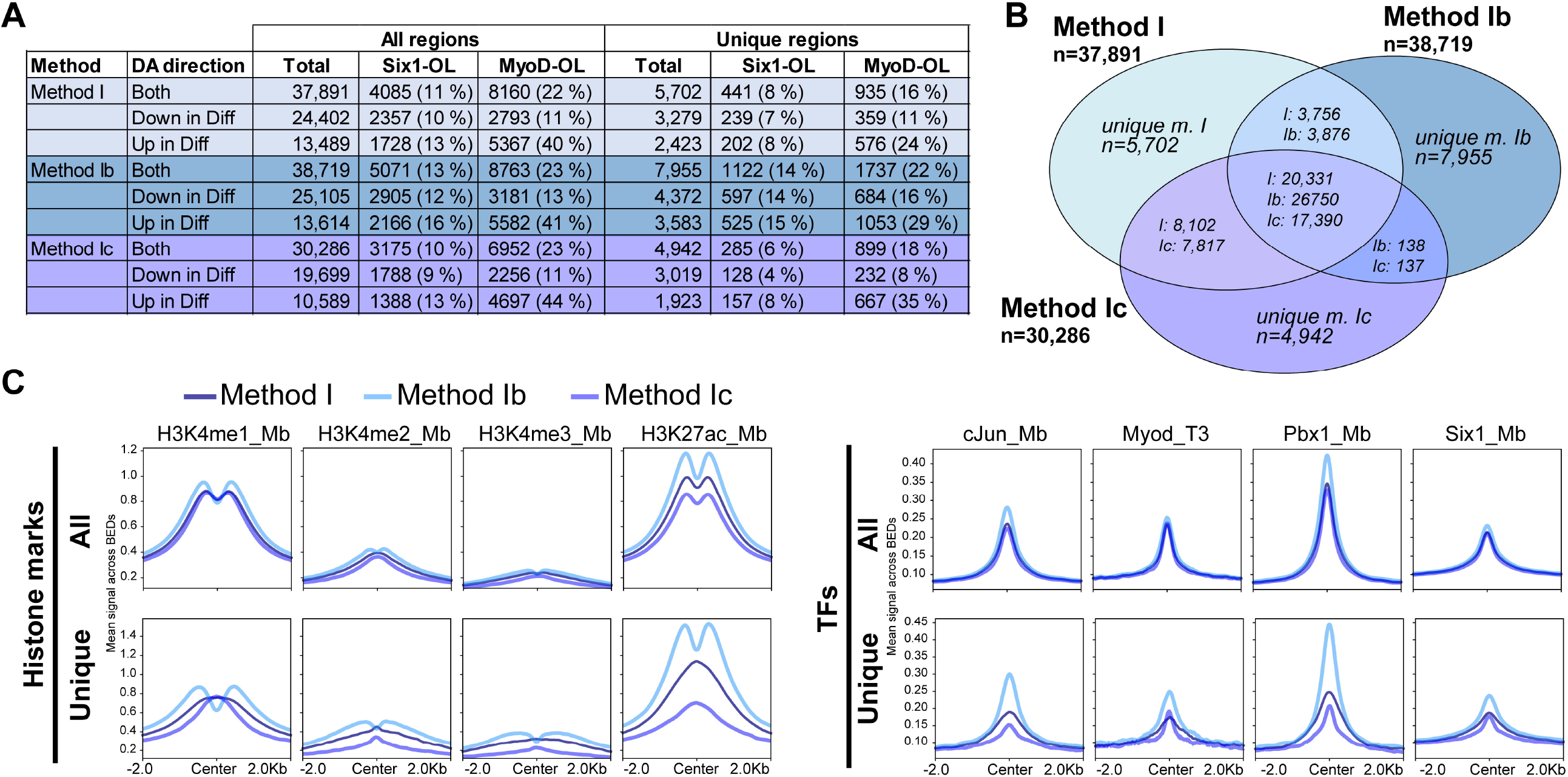
Effect of window length on sliding window approaches. Method **I** (50 bp sliding window counting of Tnp insertions) was compared with the same approach but with smaller windows (method **Ib**, 25 bp windows) or larger ones (method **Ic**, 100 bp windows). **A)** Summary of DA regions identified, broken down in those that increase or decrease in differentiating myoblasts, and with counts of overlap with Six1 or MyoD binding sites. **B)** Venn diagram showing the extent of overlap between the DA regions identified by the three methods. **C)** Mean profiles of signal from ChIP-seq experiments for histone marks and transcription factors, at the DA regions identified.

## Discussion

In this study, we have re-examined the parameters implicated in the identification of DA regions from ATAC-seq data. We have found that a one-step method that dispenses with peak calling is more sensitive. We also concluded that quantifying Tnp insertions is generally more sensitive and precise compared to methods that consider entire paired-end sequencing fragments, even if those are filtered to retain only short ones. We call this approach widaR and have included example R code that can be easily adapted to various projects.

The widaR approach has several parameters that can be adjusted by users. In particular, the length of windows, the ratio of signal in windows over genome-wide background signal, the data normalization methods, and the DA parameters (logFC and adjusted p-value for the differential test) can be adjusted based on the specifics of each experiment and the objectives of the experimenter. The availability of several normalization schemes and possibility to remove unwanted variance should make widaR suitable for the vast majority of ATAC-seq datasets. The flexibility afforded by csaw and edgeR also means that the analysis of datasets containing more than two sample types, or based on time- or dose-series, should be possible. This would represent an advantage over THOR, which only supports designs with two sample types.

An additional benefit of widaR, working with Tnp insertion sites, is that it is perfectly suitable for single-end ATAC-seq experiments. Although the majority of ATAC-seq experiments nowadays are performed in paired-end sequencing, some earlier studies were performed with single-end sequencing and could not rely on Tn5 insertion fragments. widaR performance should not suffer from such designs. Further, we note that although it was not tested in the current study, the csaw package on which widaR is based allows to use overlapping windows, which we predict should further increase the resolution and sensitivity of DA region identification.

## Methods

### Cell culture

C2C12 myoblasts were grown as described previously (*35*) in DMEM with high glucose supplemented with 2 mM glutamine and 10% fetal bovine serum. Differentiation was induced by growing cells to confluence and switching the medium to DMEM with 2 mM glutamine and 2% horse serum.

### ATAC-seq

The OMNI-ATAC protocol (*36*) was used to perform ATAC-seq on C2C12 myoblasts in proliferation phase or 24 hours following differentiation induction with 2% (v/v) horse serum. Ten thousand cells were typically used per sample. Tn5 transposomes (Tnp) were purchased from Illumina or home-made following the protocol of Picelli et al.(*37*), using pTXB1-Tn5 (gift from Rickard Sandberg, (Addgene plasmid # 60240)). Transposome were used with 2X TD buffer from the OMNI-ATAC protocol. Libraries were prepared using NEBNext Ultra II Q5 Master Mix and the set of single-index primers originally used by Buenrostro et al. (*9*). Real-time PCR on a 10% aliquot of the library after five cycles of amplification was performed to determine the total number of cycles to use to avoid over-amplification of the libraries. The libraries were purified on Zymo DNA Clean & Concentrator-5 columns and their size distributions were determined on Agilent TapeStation. After pooling, bar-coded libraries were double-sided purified with AMPure XP beads (Beckman) to remove fragments below 50 bp and above 2 kb. Libraries were sequenced on Hiseq4000 in paired-end mode at the IGBMC GenomEast genomics platform. Sequencing results are available on NCBI GEO under accession GSE195612. Basic sequencing statistics are provided in Table S1.

### ATAC-seq data processing

Reads were analyzed for quality using fastQC v0.11.9, then trimmed of adapters and low-quality sections in 3’ using fastp v0.20.1 (*38*) with the parameters --length_required 25, --cut_tail, --cut_tail_window_size 4, --cut_tail_mean_quality 20, --disable_quality_filtering. Reads were aligned in pairs to the mouse mm9 genome assembly using STAR (*39*) v2.7.5a without gene annotation GTF file, with the parameters --alignIntronMax 1 --alignEndsType Extend5pOfReads12 --alignMatesGapMax 2000, --outFilterMatchNminOverLread 0.50, --outFilterScoreMinOverLread 0.50 and filtering for properly paired mates with a mapq score of 40 or above. Duplicate reads were marked using Picard (*40*). deepTools *AlignmentSieve* (*41*) with parameter --ATACshift was used to perform shifting of the two sequencing mates by 5 and −4 nucleotides to correct for the 9 bp sequence duplication that occurs during Tnp insertion (*42*). Genome-wide sequencing coverage was calculated using deepTools v3.5.0 *bamCoverage* function. For “fragments” based analysis, where fragments represent the genomic interval between two neighboring Tnp insertion sites and are defined by the two sequencing mates, *bamCoverage* was run with the options --extendReads, --minFragLength 10 and --maxFragLength equal to either 130 (subnucleosomal fragments) or 2000 (all fragments), excluding reads marked as duplicates. To look at Tnp insertion sites independent of neighboring sites, the option --offset 1 was used, so that only coverage at the 5’ end of reads (Tnp insertion sites) would be counted. This is equivalent to what has been done in the NucleoATAC study (*15*). bigWig tracks were normalized by counts per million (CPM) in the sample. Example code to generate these files can be found in Supplementary_script.01.From_fastq_to_peaks.sh.txt. The bigWig files generated can be downloaded from the NCBI GEO entry. To simplify the bigWig tracks in example figures, the arithmetic mean of signal of replicate samples was calculated using wiggleTools v.1.2.11 (*43*). Coverage data was visualized using the IGV Browser version 2.11.9 (*44*). The Ctcf ChIP-seq dataset was obtained from GSE157339 (*45*).

ATAC-seq peaks were identified from ATAC-shifted BAM sequence files using MACS2 (*46*). When calling peaks on Tnp insertion sites, paired-end BAM files were first converted to two BED files representing each of the sequencing mates using BEDTools (*47*) before being joined back together and calling peaks in BED mode. This strategy was needed since in BAM paired-end mode MACS2 considers the entire interval delimited by the two mates, and it avoided losing information from the second mates, which are disregarded by MACS2 in BAM single-end mode (*48*). Using MACS2 settings --nomodel --shift −37 --extsize 73 essentially focused the peak calling a region of 73 base pairs centered around the Tn5 Tnp insertion site. The parameters --min-length 100 --max-gap 100 were also set, to eliminate peaks less than 100 bp in length and to merge peaks separated by less than 100 bp. When calling peaks on Tn5 Tnp intervals, MACS2 was used in BAM paired-end mode with the parameters --min-length 100 --max-gap 100. Peaks overlapping blacklisted regions were subsequently removed using BEDTools *intersectBed*. Peak calling was performed on each biological replicate separately. To arrive at a consensus set of peaks that are found in at least two replicates of a condition (growing or differentiating), rmspc (*49, 50*) was used with parameters -r bio -w 1e-5 -s 1e-10 -c 2 and using the p-value calculated by MACS2 as the ‘value’.

Summarization of Tn5 Tnp insertion sites or fragments over peaks was performed using the R package *Rsubread* and its *featureCounts* function (*51, 52*), specifying options read2pos=5 and isPairedEnd=FALSE to count all Tn5 Tnp insertions, or isPairedEnd=TRUE and checkFragLength=TRUE, minFragLength=10 and maxFragLength=130 when counting subnucleosomal fragments.

To identify DA regions with THOR, the RGT program package version 0.13.2 was used, with the following parameters: --pvalue=0.05, --exts=1,1,1,1,1,1,1, --binsize=50, --step=50 using the ATAC-shifted BAM files as input. This made the program consider only the 5’-most positions of each read with the same genomic windows used with *csaw*. The output BED-like file was imported in R/Bioconductor as a GRanges object and filtered to retain only the regions with an absolute log_2_FC value of 1 or higher.

### Sliding window analysis

The R/Bioconductor *csaw* package version 1.28.0 was used to quantify ATAC-seq data in sliding windows on ATAC-shifted BAM files. To quantify all Tnp insertions, the readParam values were set to pe=“none”, dedup=TRUE, ext=1, shift=0. Instead to quantify fragments, the parameters pe=“both”, dedup=TRUE and max.frag=1000, ext=1, shift=0 were used. To remove unwanted variation (batch effects) between the samples, the Bioconductor package RUVseq was used (*53, 54*). Inspection of RLE and PCA plots using the various algorithms (RUVg, RUVs and RUVr) determined that RUVs with k=2 was sufficient to remove the effects of batches. Significant windows were those with an absolute log_2_FC value of 1 or higher and an FDR value of 0.05 or lower. Significant DA windows were split in up- and down-regulated groups; windows in each of these groups were merged together if they were immediately adjacent (Figure S5). The R code is provided in Supplementary_script.02.widaR.R.txt.

### Algorithm performance comparisons

To determine if significant regions identified by the various methods tested are biologically significant, three methods were used. Assignment of genes to regions was done with the package *ChIPpeakanno* (*55*), taking the closest gene within a maximum distance of 50 kb. GO biological process term enrichment for the genes associated to the regions was performed in R/Bioconductor (*56*) with the *topGO* package (*57*). Enrichment was calculated as the ratio of observed over expected number of genes in each category, considering all genes in the mouse genome as background. A Benjamini-Hochberg adjusted p value of 1E-4 was the cut-off for significance. Enrichment of epigenetic marks (H3K4me1, H3K4me3 and H3K27ac (*58, 59*)) and transcription factors (c-Jun in C2C12 myoblasts from GSE37525 (*59*), Pbx1 in C2C12 myoblasts from GSE76010 (*60*), Six1 in primary myoblasts from GSE175999 (*61*), MyoD at 3 hours of differentiation in C2C12 from GSE80812 (*62*)) associated with gene regulatory elements in muscle precursors was evaluated using deepTools *computeMatrix* and *plotHeatmap*. The arithmetic mean signal at all regions found by a given method, or all regions uniquely found by a given method, was plotted. The median signal was also examined to determine if outliers were skewing the mean values, and qualitatively similar results were obtained (Figure S6). mRNA expression levels of the genes closest to the DA regions was analyzed, using a dataset consisting of five growth phase C2C12 samples and two samples at 24 hours of differentiation (to be published elsewhere). RNA was prepared by Trizol extraction and DNAse digestion, and sequencing libraries were prepared using the TrueSeq kit (Illumina), using ribosomal RNA depletion. Sequencing was done in paired-end layout on a HiSeq4000. Sequence data were analyzed using a standard pipeline, including read trimming and alignment with STAR, summarization over ENSEMBL mouse genes using featureCounts, removeal of low-count genes, normalization using TMM and RUVseq, and differential expression testing with edgeR and glmQLFTest function using a design to contrast differentiated cells with proliferating myoblasts. The results of differential expression testing are provided in supplementary dataset 1 (Supplementary_dataset_01.RNAseq_differential_expression_C2C12.txt).

## Supporting information

Supplementary_script.01

Supplementary_script.02

## Declarations

### Ethics approval and consent to participate

Not applicable.

### Consent for publication

Not applicable.

### Availability of data and materials

The ATAC-seq data for C2C12 myoblast differentiation has been deposited on the NCBI Gene Expression Omnibus (GEO) under accession GSE195612.

### Authors’ contributions

AB designed the study, secured funding, performed ATAC-seq experiments, developed the algorithm, analyzed data, and wrote the manuscript. AS developed the algorithm, analyzed data, and wrote the manuscript. All authors read, edited, and approved the final manuscript.

## Acknowledgements

The authors would like to thank the following individuals or groups for their contributions: members of the Blais lab (uOttawa) and members of the team of Irwin Davidson (IGBMC) for helpful discussions; Isabelle Michel (IGBMC) for technical support; the bioinformatics group of the IGBMC for helpful discussions related to data analysis; the GenomEast and tissue culture facility staff of the IGBMC; Compute Canada for access to the Cedar high-performance computing cluster. The authors acknowledge the financial support of the Canadian Institutes of Health Research and the Chaires de Recherche Gutenberg program.

## Figures and legends

**Figure S1. Number of DNA accessibility mapping studies reported in PubMed, separated by method**.

A PubMed search with the indicated terms was performed on February 1, 2022. The number of resulting articles was per year is indicated as a fraction of all articles mentioning at least one of the three methods. Articles mentioning more than one method are counted in each relevant group.

**Figure S2. ATAC-seq peak calls**.

**A-F)** Distribution of length (genomic width, in base pairs) of the peaks identified using MACS2 on ATAC-shifted sequencing fragments in growing or differentiating cells, or on the union of both peak sets. A-C show length of peaks when they are called using sequencing fragments (BAMPE mode in MACS2) while D-E show length on peaks called using transposase insertion sites. **G-I)** Overlap between peaks called using fragments or insertions, in growing or differentiating, or the union of both peak sets. The overlap was calculated in a base-wise fashion and is reported in kilobase of overlapping peak regions, to account for partially overlapping peaks. **J)** Overlap between peaks called using fragments or insertions, identified at the peak level. The intersection part was expressed in terms of peaks from each set, because some peaks found using one method overlap two peaks found with the other method. **K-M)** Overlap between peaks identified in growing and in differentiating cells. Shown using base-wise overlap (J, K) or region-wise overlap (L, M), on peaks called with fragments (J, L) or insertion sites (K, M). Based on these results,

**Figure S3. Comparison of TMM and LOESS normalization methods on sliding-window data**.

RLE and PCA plot were prepared from the data using methods I and II, applying either TMM on high-signal windows (5-fold more signal than on genome-wide background) or LOESS (as implemented by the normOffsets command of csaw). Additionally, a DA test was performed comparing Diff and Grow samples. MA plots showing the logFC between the two sample types as a function of the signal across samples are shown. The blue lines show the data trend with a LOESS fit while the green dotted lines show the cutoff for abs(log_2_FC) > 1. Red circles represent windows meeting the logFC criterion and having FDR values smaller than 0.05; their numbers are indicated in the upper and lower sections of the MA plots.

**Figure S4. Effect of batch effect removal with RUVseq**.

Method I data were TMM normalized, and optionally batch effects were removed using RUVr or RUVs algorithms, with different values of k. With RUVr, all windows were used as invariant set. Similar results were obtained with methods II and III (not shown).

**Figure S5. Strategy used to merge significant windows into regions**.

When sliding window-based methods are used, neighboring windows that significantly change are merged together into significant regions. This only occurs if windows are immediately adjacent, and if the change occurs in the same direction (greater or lower DA in condition 2 versus 1). For diagram simplicity, only log_2_ fold-change is shown as determinant of DA significance.

**Figure S6. Median and mean signal give comparable results**.

Related to Figure 3. Plots representing the average signal for genomic features (presence of histone modification marks or binding of transcription factors) are generated to obtain an estimate of how the different methods perform in identifying biologically meaningful DA regions. Using the arithmetic mean or the median of the signal at the regions found by each method yields results that are qualitatively similar.

**Figure S7. Heatmaps and mean plots for the comparison of methods I, II and III**.

Related to Figure 3. Full analysis of histone mark and transcription factor binding ChIP-seq signal, including heatmaps.

**Figure S8. Effect of calling accessible DNA peaks using sequenced fragments or insertion sites**.

MACS2 was used to call peaks on entire sequenced fragments (BAMPE mode) or with segments centered on the 5’ ends of each read (mates) of pairs (BED mode). The mean signal profiles of histone modification marks and TF binding are all superior when insertion peaks are used (light colors, compared to darker shades). Additionally, both methods perform best when used for the quantification of ATAC-seq fragments limited to those that are of sub-nucleosomal length instead of fragments of all lengths (green, compared to blue).

**Figure S9. Length distribution of DA regions, for each method**.

The genomic length (in base pair) distribution for each analysis method. For sliding window-based methods, this is done after merging adjacent significant windows.

## Other supplementary material

***Supplementary_script.01.From_fastq_to_peaks.sh.txt***

Shell script for the analysis of ATAC-seq datasets, starting from fastq files, and until the generation of peaks for each individual replicate.

***Supplementary_script.02.widaR.R.txt***

R script for the identification of DA regions by a sliding window approach, starting from ATAC-seq BAM files.

***Supplementary_dataset_01.RNAseq_differential_expression_C2C12.txt***

RNA-seq expression data with differential expression test results, for the contrast “Diff - Grow”. Only genes with at least minimum expression counts of 10 in at least 2 samples were retained.

