## Supplementary_script.01 for "Improved sensitivity and resolution of ATAC-seq differential DNA accessibility analysis"

################################################################################
################################################################################
### Handling input files and running with SLURM job scheduler
# An "sbatch" script passes every input file individually to a "launcher" script, along with sample information
# The launcher script sets the SLURM variables and launches the "executable" script that follows below

################################################################################
### sbatch script (move this to a separate file)
#!/bin/bash
files='ATAC_C2C12_Grow_r1.LRBS1 ATAC_C2C12_Grow_r2.LRBS9 ATAC_C2C12_Grow_r3.LRBS16 ATAC_C2C12_Grow_r4.LRBS52 ATAC_C2C12_Diff_r1.LRBS5 ATAC_C2C12_Diff_r2.LRBS27 ATAC_C2C12_Diff_r3.LRBS65'
# Use a loop, where the same code is executed for each element of the list above
for data in $files
do
# Period is set as delimiter
IFS='.'
# Split information
read -ra ADDR <<< "$data" 
IFS=' '
sample_name=${ADDR[0]}
# Send to launcher script, along with the file relevant information
sbatch -J $sample_name.%j -o $sample_name.%j.out --export=VARIABLE=$data My_Launcher_Script.sh
done
################################################################################

################################################################################
### My_Launcher_Script.sh (move this to a separate file)
#!/bin/bash
#SBATCH -n 16
#SBATCH -N 1
#SBATCH --mem 45G
#SBATCH -t 1-0:00
./My_Executable_script.sh ${VARIABLE}
################################################################################

################################################################################
### My_Executable_script.sh (from here and all the way to the end of this file) (move this to a separate file)
#!/bin/bash

# Input file info received from My_Launcher_Script
i=$1

### Keep track of execution time
# This records the time when the execution started
res1=$(date +%s.%N)

########################################
### ATAC-seq analysis , steps performed:
# -QC analysis with fastqc, both before and after read trimming
# https://www.bioinformatics.babraham.ac.uk/projects/fastqc/
# -Merge reads coming from each lane of the sequencing flow cell
# -QC and trimming of reads using fastp
# https://github.com/OpenGene/fastp
# -alignment on genome using STAR, index without GTF and max intron length of 1 bp
# https://github.com/alexdobin/STAR
# -indexing of BAM file with samtools
# http://www.htslib.org/doc/samtools.html
# -marking duplicates with Picard
# https://broadinstitute.github.io/picard/command-line-overview.html#MarkDuplicates
# -Adjusts BAM files by ATAC shifting
# https://deeptools.readthedocs.io/en/develop/content/tools/alignmentSieve.html
# -Create tracks representing read density along the genome, with deepTools bamCoverage
# https://deeptools.readthedocs.io/en/develop/content/tools/bamCoverage.html
# -Call peaks using MACS2 in BED and BAMPE modes
# https://github.com/macs3-project/MACS


########################################
### Create variables that will be used throughout the script
### Adjust according to own system

# Path to where home projects directory is
home_projects=$HOME/my_projects/
# Give a name to this specific project
project_name=My_project_name
# define temporary project directory, on scratch for fast data access and computation
project_dir=$HOME/scratch/$project_name
# Temporary file folder used by certain processes
temp_directory=$HOME/scratch/temp
# Location of original fastq files received from sequencing facility
ori_fastq=$project_dir/fastq_files
# Genome information directory
genome_dir=$home_projects/Genomes/mm9/
# Genome blacklist
blacklist=$genome_dir/mm9-blacklist.bed
# Genome index for STAR alignment
star_index=$genome_dir/star_index_noAnnot_v2.7.5a
# File name extension to give to the STAR output
starFileExt=star_index_noAnnot
# Genome for MACS2
macs_organism=mm


########################################
### Create directories to receive the output files
### To be done only once ahead of running the script

# # Define project name
# project_name=My_project_name
# # make directory for entire project
# mkdir $HOME/scratch/$project_name
# # define temporary project directory on scratch
# project_dir=$HOME/scratch/$project_name
# # make directory to receive fastQC reports
# mkdir $project_dir/fastQC_reports
# # make directory to receive fastp reports
# mkdir $project_dir/fastp_reports
# # make directory to receive fastq files 
# mkdir $project_dir/fastq_files
# # make directory to receive STAR output bam files 
# mkdir $project_dir/bam_and_bai_files
# # make directory to receive STAR log files 
# mkdir $project_dir/STAR_reports
# # make directory to receive Picard report files 
# mkdir $project_dir/picard_reports
# # make directory to receive deepTools bigWig density files 
# mkdir $project_dir/bedgraph_bigwig_files
# # make directory to receive MACS2 peak calling output
# mkdir $project_dir/peak_calling_MACS2

#######################################
### File name split to get the sample name and accession number
IFS='.' 
read -ra ADDR <<< "$i" 
IFS=' '

# assign new variable names to the substrings
acc=${ADDR[1]}
sample_name=${ADDR[0]}

# Save information about computation progress in : process_report_file
process_report_file=$project_dir/$sample_name.process_report.out

echo "##################################################################" >>$process_report_file
echo "##################################################################" >>$process_report_file
echo "##################################################################" >>$process_report_file
echo Working with:		$acc >>$process_report_file
echo Sample name:		$sample_name >>$process_report_file


#####################################################
#####################################################
### Start of computation

### load required modules
module load StdEnv/2020
module load fastp/0.20.1 star/2.7.5a picard/2.23.3 samtools/1.11 fastqc/0.11.9 bedtools/2.29.2

echo "#################################" >>$process_report_file
echo "#################################" >>$process_report_file
echo fastQC prior to trimming for $i >>$process_report_file

### Analysis of original fastq files
cat $ori_fastq/$acc.R1.fastq.gz $ori_fastq/$acc.R2.fastq.gz |\
fastqc stdin:$sample_name.pre_trimming \
-o $project_dir/fastQC_reports/ \
-f fastq \
--contaminants $home_projects/Genomes/fastQC_sequences/contaminant_list.txt \
--adapters $home_projects/Genomes/fastQC_sequences/adapter_list.txt \
--limits $home_projects/Genomes/fastQC_sequences/limits.txt \
-d $temp_directory \
>>$process_report_file 2>&1


#####################################################
#####################################################
### Use fastp to remove low quality sections of reads and remove sequencing adapter contamination

echo "#################################" >>$process_report_file
echo "#################################" >>$process_report_file
echo fastp for $i >>$process_report_file

fastp \
--in1 $project_dir/fastq_files/$acc.R1.fastq.gz \
--in2 $project_dir/fastq_files/$acc.R2.fastq.gz \
--thread 16 \
--length_required 25 \
--adapter_sequence CTGTCTCTTATACACATCTCCGAGCCCACGAGAC \
--adapter_sequence_r2 CTGTCTCTTATACACATCTGACGCTGCCGACGA \
--cut_tail \
--cut_tail_window_size 4 \
--cut_tail_mean_quality 20 \
--disable_quality_filtering \
--overrepresentation_analysis \
--overrepresentation_sampling 10 \
--json $project_dir/fastp_reports/$sample_name.fastp_report.json \
--html $project_dir/fastp_reports/$sample_name.fastp_report.html \
--out1 $project_dir/fastq_files/$sample_name.trimmed.R1.fastq \
--out2 $project_dir/fastq_files/$sample_name.trimmed.R2.fastq \
>>$process_report_file 2>&1


echo "#################################" >>$process_report_file
echo "#################################" >>$process_report_file
echo fastQC after trimming for $i >>$process_report_file

### Analysis of fastq R1+R2 file quality, after trimming
cat $project_dir/fastq_files/$sample_name.trimmed.R1.fastq $project_dir/fastq_files/$sample_name.trimmed.R2.fastq |\
fastqc \
stdin:$i.trimmed \
-o $project_dir/fastQC_reports/ \
-f fastq \
--contaminants $home_projects/Genomes/fastQC_sequences/contaminant_list.txt \
--adapters $home_projects/Genomes/fastQC_sequences/adapter_list.txt \
--limits $home_projects/Genomes/fastQC_sequences/limits.txt \
-d $temp_directory \
>>$process_report_file 2>&1


#####################################################
#####################################################
### Use STAR to map reads on the genome (i.e. find where the read sequences came from)
### Done with STAR
# For ATAC-seq, a genome index created without specifying a gene structure GTF file is used
# Additional criteria relevant to ATAC-seq: maximum INTRON size is 1 bp
# Additional criteria relevant to ATAC-seq: maximum FRAGMENT size is 2000 bp
# Additional criteria relevant to ATAC-seq: align with Extend5pOfReads12 (soft-clipping possible only at the 3'end) 
#	...the 5' end of reads MUST align perfectly to reference genome without soft-clipping
#	...this is important since we quantitate ATAC signal by looking at insertions, which occur at the 5' end
echo "#################################" >>$process_report_file
echo "#################################" >>$process_report_file
echo STAR for $i >>$process_report_file

STAR \
--runThreadN 16 \
--alignIntronMax 1 \
--alignEndsType Extend5pOfReads12 \
--alignMatesGapMax 2000 \
--outFilterMatchNminOverLread 0.50 \
--outFilterScoreMinOverLread 0.50 \
--genomeDir $star_index \
--readFilesIn $project_dir/fastq_files/$sample_name.trimmed.R1.fastq $project_dir/fastq_files/$sample_name.trimmed.R2.fastq \
--outSAMtype BAM SortedByCoordinate \
--outSAMmapqUnique 40 \
--outFilterMultimapNmax 1 \
--outFileNamePrefix $project_dir/bam_and_bai_files/$sample_name.$starFileExt. \
>>$process_report_file 2>&1

# Move report files created by STAR to dedicated report directory
mv $project_dir/bam_and_bai_files/$sample_name.$starFileExt.Log.out $project_dir/STAR_reports/
mv $project_dir/bam_and_bai_files/$sample_name.$starFileExt.Log.final.out $project_dir/STAR_reports/
mv $project_dir/bam_and_bai_files/$sample_name.$starFileExt.Log.progress.out $project_dir/STAR_reports/

# Index the BAM file made by STAR
echo "#################################" >>$process_report_file
echo "#################################" >>$process_report_file
echo Indexing after STAR for $i >>$process_report_file

samtools index -@ 16 $project_dir/bam_and_bai_files/$sample_name.$starFileExt.Aligned.sortedByCoord.out.bam \
$project_dir/bam_and_bai_files/$sample_name.$starFileExt.Aligned.sortedByCoord.out.bam.bai \
>>$process_report_file 2>&1


#####################################################
#####################################################
### Identify read pairs that are duplicates caused by PCR amplification
echo "#################################" >>$process_report_file
echo "#################################" >>$process_report_file
echo Deduping $i >>$process_report_file

java -jar $EBROOTPICARD/picard.jar MarkDuplicates \
-VERBOSITY WARNING \
-I $project_dir/bam_and_bai_files/$sample_name.$starFileExt.Aligned.sortedByCoord.out.bam \
-O $project_dir/bam_and_bai_files/$sample_name.$starFileExt.markedDups.bam \
-M $project_dir/picard_reports/$sample_name.$starFileExt.marked_dup_metrics.txt \
-TMP_DIR $temp_directory \
>>$process_report_file 2>&1

### Index the new BAM file made with marked duplicates
echo "#################################" >>$process_report_file
echo "#################################" >>$process_report_file
echo indexing after Picard for $sample_name >>$process_report_file

samtools index -@ 16 $project_dir/bam_and_bai_files/$sample_name.$starFileExt.markedDups.bam \
$project_dir/bam_and_bai_files/$sample_name.$starFileExt.markedDups.bam.bai \
>>$process_report_file 2>&1


#####################################################
#####################################################
### Collect statistics on the distribution of insert sizes in the ATAC-seq experiment
# load required modules
module load r/4.0.2 gcc/9.3.0
module load gsl/2.6

echo "#################################" >>$process_report_file
echo "#################################" >>$process_report_file
echo Picard CollectInsertSizeMetrics for $sample_name >>$process_report_file

java -jar $EBROOTPICARD/picard.jar CollectInsertSizeMetrics \
-I $project_dir/bam_and_bai_files/$sample_name.$starFileExt.markedDups.bam \
-O $project_dir/picard_reports/$sample_name.$starFileExt.markedDups.insert_metrics_table.txt \
-H $project_dir/picard_reports/$sample_name.$starFileExt.markedDups.insert_metrics_histogram.pdf \
-HISTOGRAM_WIDTH 2000 \
-VERBOSITY WARNING \
>>$process_report_file 2>&1


#####################################################
#####################################################
### Run deepTools alignmentSieve to perform ATACshift on the BAM files
# deepTools is run from a virtual environment
module load python/3.8.2
module load scipy-stack/2020b
# Activate the deepTools python environment
source $home_projects/env_DEEPTOOLS/bin/activate

echo "#################################" >>$process_report_file
echo "#################################" >>$process_report_file
echo alignmentSieve on $i >>$process_report_file

# Perform ATACshifting of the BAM file
alignmentSieve \
--numberOfProcessors 16 \
--bam $project_dir/bam_and_bai_files/$sample_name.$starFileExt.markedDups.bam \
--ATACshift \
--outFile $project_dir/bam_and_bai_files/$sample_name.$starFileExt.markedDups.ATACshiftpresort.bam \
>>$process_report_file 2>&1

# Sort the new ATACshifted BAM file
samtools sort -@ 16 $project_dir/bam_and_bai_files/$sample_name.$starFileExt.markedDups.ATACshiftpresort.bam \
-o $project_dir/bam_and_bai_files/$sample_name.$starFileExt.markedDups.ATACshift.bam \
>>$process_report_file 2>&1

# Index the new ATACshifted BAM file
samtools index $project_dir/bam_and_bai_files/$sample_name.$starFileExt.markedDups.ATACshift.bam \
$project_dir/bam_and_bai_files/$sample_name.$starFileExt.markedDups.ATACshift.bam.bai \
>>$process_report_file 2>&1


#####################################################
#####################################################
### Create read density (genome coverage) graphs using deepTools
echo "#################################" >>$process_report_file
echo "#################################" >>$process_report_file
echo bamCoverage 10-130 bp on $i >>$process_report_file

# Here coverage is calculated only using fragments that certainly come from "nucleosome-free" regions 
#    due to the fragment length being between 10 and 130 bp, shorter than the length of one nucleosome (~147 bp)

bamCoverage \
--numberOfProcessors 16 \
--outFileFormat bigwig \
--normalizeUsing CPM \
--binSize 10 \
--minFragmentLength 10 \
--maxFragmentLength 130 \
--ignoreDuplicates \
--bam $project_dir/bam_and_bai_files/$sample_name.$starFileExt.markedDups.ATACshift.bam \
--ignoreForNormalization chrX chrM \
--blackListFileName $blacklist \
--outFileName $project_dir/bedgraph_bigwig_files/$sample_name.$starFileExt.ignoreDups.ATACshift10-130bin10CPM.bw \
>>$process_report_file 2>&1

echo "#################################" >>$process_report_file
echo "#################################" >>$process_report_file
echo bamCoverage 10-2000 bp on $i >>$process_report_file

# Here coverage is calculated only using fragments that certainly come from "nucleosome-free" regions 
#    due to the fragment length being between 10 and 130 bp, shorter than the length of one nucleosome (~147 bp)

bamCoverage \
--numberOfProcessors 16 \
--outFileFormat bigwig \
--normalizeUsing CPM \
--binSize 10 \
--minFragmentLength 10 \
--maxFragmentLength 2000 \
--ignoreDuplicates \
--bam $project_dir/bam_and_bai_files/$sample_name.$starFileExt.markedDups.ATACshift.bam \
--ignoreForNormalization chrX chrM \
--blackListFileName $blacklist \
--outFileName $project_dir/bedgraph_bigwig_files/$sample_name.$starFileExt.ignoreDups.ATACshift10-2000bin10CPM.bw \
>>$process_report_file 2>&1


# Here instead coverage is calculated at sites of Tn5 insertion 
#     Each end of fragment represents an insertion, this is set with the --Offset 1 option
echo "#################################" >>$process_report_file
echo "#################################" >>$process_report_file
echo bamCoverage Offset 1 on $i >>$process_report_file

bamCoverage \
--numberOfProcessors 16 \
--outFileFormat bigwig \
--normalizeUsing CPM \
--Offset 1 \
--binSize 10 \
--ignoreDuplicates \
--bam $project_dir/bam_and_bai_files/$sample_name.$starFileExt.markedDups.ATACshift.bam \
--ignoreForNormalization chrX chrM \
--blackListFileName $blacklist \
--outFileName $project_dir/bedgraph_bigwig_files/$sample_name.$starFileExt.ignoreDups.ATACshiftOffset1bin10CPM.bw \
>>$process_report_file 2>&1


#####################################################
#####################################################
### Identify "peaks" of ATAC-seq signal (sites where Tn5 insertion is frequent)

echo "#################################" >>$process_report_file
echo "#################################" >>$process_report_file
echo Preparing BAM files for MACS2 in BED format for $i >>$process_report_file

# prepare the BAM files to be analyzed by MACS2 by converting them to BED format
# Strategy is to convert BAM to BED and to split the two mates into separate files
#	...then to bring them together again by providing both as input in the 'macs2 callpeak' command
#	...further, we run MACS2 with --shift and --extsize so that in reality the reads become centered at the insertion site 
#	...the result is that we end up calling peaks on 100% of the info, and on insertions instead of fragments

# samtools with -f 64 restricts the output to only those reads (mates) that are FIRST IN PAIR
# note that 64 will RETAIN those reads marked as duplicates. 
# If it is desired to EXCLUDE them, the following could be used instead: samtools view -hf 64 - | samtools view -hF 1024 -
samtools view -hf 64 $project_dir/bam_and_bai_files/$sample_name.$starFileExt.markedDups.ATACshift.bam |\
bedtools bamtobed -i stdin >$project_dir/bam_and_bai_files/$sample_name.$starFileExt.markedDups.ATACshift.first.bed

# samtools with -f 128 restricts the output to only those reads (mates) that are SECOND IN PAIR
# note that 128 will RETAIN those reads marked as duplicates. 
# If it is desired to EXCLUDE them, the following could be used instead: samtools view -hf 128 - | samtools view -hF 1024 -
samtools view -hf 128 $project_dir/bam_and_bai_files/$sample_name.$starFileExt.markedDups.ATACshift.bam |\
bedtools bamtobed -i stdin >$project_dir/bam_and_bai_files/$sample_name.$starFileExt.markedDups.ATACshift.second.bed


# Load modules for MACS2 peak caller
deactivate
module purge
module load StdEnv/2020
module load python/3.8.2 bedtools/2.29.2
# activate the MACS2 python environment
source $home_projects/env_MACS2/bin/activate

# Run MACS2 in BED mode, specifying both reads on the input line
echo "#################################" >>$process_report_file
echo "#################################" >>$process_report_file
echo MACS2 in BED format on $i >>$process_report_file

macs2 callpeak \
-t $project_dir/bam_and_bai_files/$sample_name.$starFileExt.markedDups.ATACshift.first.bed \
$project_dir/bam_and_bai_files/$sample_name.$starFileExt.markedDups.ATACshift.second.bed \
-f BED \
-g mm \
--shift -37 --extsize 73 \
--nomodel --keep-dup all \
--min-length 100 --max-gap 100 \
--outdir $project_dir/peak_calling_MACS2/ \
--name $sample_name.MACS2_bed_atacshift_wDups_s37_e73_min100max100 \
>>$process_report_file 2>&1


# Then run MACS2 in BAMPE mode.
echo "#################################" >>$process_report_file
echo "#################################" >>$process_report_file
echo MACS2 in BAMPE format on $i >>$process_report_file

macs2 callpeak \
-t $project_dir/bam_and_bai_files/$sample_name.$starFileExt.markedDups.ATACshift.bam \
-f BAMPE \
-g mm \
--nomodel --keep-dup all \
--min-length 100 --max-gap 100 \
--outdir $project_dir/peak_calling_MACS2/ \
--name $sample_name.MACS2_bampe_atacshift_wDups_min100max100 \
>>$process_report_file 2>&1

# Remove blacklisted regions from MACS2 output narrowPeak files
echo "#################################" >>$process_report_file
echo "#################################" >>$process_report_file
echo Removing blacklisted regions from peaks on $i >>$process_report_file

bedtools intersect -v \
-a $project_dir/peak_calling_MACS2/$sample_name.MACS2_bed_atacshift_wDups_s37_e73_min100max100_peaks.narrowPeak \
-b $blacklist \
>$project_dir/peak_calling_MACS2/$sample_name.MACS2_bed_atacshift_wDups_s37_e73_min100max100.noBL.narrowPeak

bedtools intersect -v \
-a $project_dir/peak_calling_MACS2/$sample_name.MACS2_bampe_atacshift_wDups_min100max100_peaks.narrowPeak \
-b $blacklist \
>$project_dir/peak_calling_MACS2/$sample_name.MACS2_bampe_atacshift_wDups_min100max100.noBL.narrowPeak


#####################################################
#####################################################
### Keep track of execution time
# This records the time when the execution ended
res2=$(date +%s.%N)

# This calcualtes the difference in time between start and end
dt=$(echo "$res2 - $res1" | bc)
dd=$(echo "$dt/86400" | bc)
dt2=$(echo "$dt-86400*$dd" | bc)
dh=$(echo "$dt2/3600" | bc)
dt3=$(echo "$dt2-3600*$dh" | bc)
dm=$(echo "$dt3/60" | bc)
ds=$(echo "$dt3-60*$dm" | bc)

### Calculate time difference and report it at the end.
echo "#################################" >>$process_report_file
echo Subject: Finished $sample_name >>$process_report_file
printf "Total runtime (days-hours-minutes-seconds-nanosec): %d:%02d:%02d:%02.4f\n" $dd $dh $dm $ds >>$process_report_file
