## Supplementary_script.02 for "Improved sensitivity and resolution of ATAC-seq differential DNA accessibility analysis"

##### ATACSeq differential accessibility analysis with CSAW #####
# Inspired by the CSAW handbook: http://bioconductor.org/books/3.15/csawBook/

##### Setup R environment ##### 
# Set path to working directory that contains data files required for analysis
setwd("~/path/to/working/directory")

# Load libraries essential for analysis
library(BiocParallel) # Enables parallelization
library(csaw) # Used for window based analysis
library(edgeR) # Used for differential testing
library(rtracklayer) # Used to import BED files as GRanges or export GRanges as BED files
library(RUVSeq) # Used for normalization, remove unwanted variance

# Experiment specific libraries
library(BSgenome.Mmusculus.UCSC.mm9) # Genome sequence for mm9, use respective library for genome being analyzed, ex: hg38, hg19, etc.

# Genome being studied
my_bs_genome <- BSgenome.Mmusculus.UCSC.mm9

# Give a name to the analysis being performed
analysis <- "My_ATAC_project" # this is done for later convenience

# Specify directory to export results to
exportDir <- "results/bedExports/"

# Create export directories in case they do not exist
dir.create(exportDir, showWarnings = TRUE, recursive = TRUE)

# Create a BiocParallel object that uses specified number of CPU threads with a progress bar
my_multicoreParam <- MulticoreParam(workers = 12, progressbar = TRUE, exportglobals = FALSE)
register(my_multicoreParam)

# Load tab-delimited .txt file with information on ATACSeq samples (i.e, name, treatment, drug, and the path to appropriate bam file)
	# ATAC-shifted BAM files are used.
	# See bottom of this file for an example of the expected structure
atac_samples <- as.data.frame(read.table("My_ATAC_Files_Shifted.txt", sep="\t", header=TRUE))
# Set sample names as row names for later downstream analysis
rownames(atac_samples) <- atac_samples$Sample
# Specify path to bam files of ATAC samples
bam.files <- paste(atac_samples$BAM_Path)
# Design to compare samples, matching order of BAM files
	# Adjust as needed depending on the study design
treatment <- factor(atac_samples$Group)
design <- model.matrix(~0 + treatment)
colnames(design) <- levels(treatment)
treat1.vs.treat2 <- makeContrasts(treat1 - treat2, levels = design )

# Specify chromosomes to limit analysis to
# Optionally can limit to a specific chromosome, ex: chrLimit <- "chr19"
	# adjust as needed based on genome being studied
chrLimit <- paste0("chr", c(1:19, "X", "Y"))
# Set regions to discard by importing blacklist loci BED file
	# adjust as needed depending on genome being studied
blacklistPath <- "bed_files/mm9-blacklist.bed"

##### CSAW-specific parameters
# Set parameters for CSAW on regions to discard and chromosomes to restrict analysis to, indicate if data is paired-end
# To count each read end independently (Tnp insertions), set 'pe="none"'
my_pe_mode <- "none"
# indicate whether to remove duplicates (TRUE) or not (FALSE)
duplicate_removal <- TRUE
# Set Window Size
windowSize <- 50
# Set spacing between windows. Use 'windowSize' to have adjacent windows without intervening gaps
windowSpacing <- windowSize
# Set the minimum number of counts for a window to be retained 
minCount_filterSize <- 50
# Set bin size for calculating global enrichment
bin.size <- 10000
# Set the fold change value to filter windows for enrichment over global enrichment in the binned dataset
globalBckg_foldChange <- 5
##### No further parameters to adjust #####


##### CSAW Analysis #####
blacklist <- import.bed(blacklistPath)
insertParam <- readParam(discard=blacklist, restrict=chrLimit, pe=my_pe_mode, dedup=duplicate_removal, minq=NA)

# Count genome into bins for filtering via fold change over global background using the insertParam
binned.data <- windowCounts(bam.files, width=bin.size, bin=TRUE, param=insertParam, BPPARAM=my_multicoreParam)

# Count Tn5 integration sites with CSAW
window.data <- windowCounts(bam.files, spacing=windowSpacing, width=windowSize, ext=1, shift=0, filter=minCount_filterSize, bin=FALSE, param=insertParam, BPPARAM = my_multicoreParam)

# Filter data by fold change
# Create a filter of windows that have signal well above global background
keep.fcFilter <- filterWindowsGlobal(window.data, background = binned.data)$filter > log2(globalBckg_foldChange)
summary(keep.fcFilter)
# Filter out unwanted windows by fold change
filtered.winData <- window.data[keep.fcFilter,]
# Calculate normalization factors on filtered high-abundance windows with CSAWs implementation of the TMM method to remove efficiency biases. 
filtered.winData <- normFactors(filtered.winData)

# Differential Analysis
# Create an EdgeR object from our normalized data, set appropriate column and row names
y.TMM <- asDGEList(filtered.winData)
colnames(y.TMM$counts) <- atac_samples$Sample
rownames(y.TMM$samples) <- atac_samples$Sample
# Estimate dispersions and fit a GLM to data
y.TMM <- estimateDisp(y.TMM, design)
fit.TMM <- glmQLFit(y.TMM, design, robust=TRUE)
# Generate a SeqExpressionSet from TMM data to use with RUVs
set.TMM <- newSeqExpressionSet(counts=y.TMM$counts, phenoData=atac_samples)
cpm.TMM <- cpm(y.TMM, normalized.lib.sizes = TRUE, log=FALSE)
multiplier.TMM <- 1e-6*(mean(y.TMM$samples$lib.size))
normCounts(set.TMM) <- round(multiplier.TMM*cpm.TMM, digits=0)

# Define the replicate groups
replicateGroups <- makeGroups(atac_samples$Group)
replicateGroups
# Create a RUVs set for our data. RUVr or RUVg may instead be used
set.RUVs <- RUVs(set.TMM, rownames(set.TMM), k=2, replicateGroups)
# Create a design matrix for differential testing that incorporates W, the variable representing unwanted variance. 
design.RUVs <- model.matrix(~0 + treatment + W_1 + W_2, data = pData(set.RUVs))
# Make contrasts for RUVs data based on the new design matrix
treat1.vs.treat2.RUVs <- makeContrasts(treatmenttreat1 - treatmenttreat2, levels=design.RUVs)

# Estimate dispersion with one command
y.RUVs <- estimateDisp(y.TMM, design.RUVs)
# Fit QL model to data
fit.RUVs <- glmQLFit(y.RUVs, design.RUVs)
# Test for differential gene expression
results.RUVs <- glmQLFTest(fit.RUVs, contrast=treat1.vs.treat2.RUVs)
# Apply FDR to results table
resultsTable.RUVs <- topTags(results.RUVs, adjust.method="BH", n=Inf, sort.by="none")$table
# Calculate scores using the FDR values
resultsTable.RUVs$score <- -10*log10(resultsTable.RUVs$FDR)
# Add FDR corrected results data to our filtered dataset of windows
winResults.Data <- filtered.winData
rowData(winResults.Data) <- cbind(rowData(winResults.Data), resultsTable.RUVs)

# Create RSE objects subsetted for significant windows
winResults.Sig.All <- subset(winResults.Data, abs(logFC) >= 1 & FDR < 0.05)
winResults.Sig.Up <- subset(winResults.Data, logFC >= 1 & FDR < 0.05)
winResults.Sig.Down <- subset(winResults.Data, logFC <= -1 & FDR < 0.05)
# Isolate GRanges objects from the RSE
window.GR.Sig.All <- rowRanges(winResults.Sig.All)
window.GR.Sig.Up <- rowRanges(winResults.Sig.Up)
window.GR.Sig.Down <- rowRanges(winResults.Sig.Down)
# Add scores to the GRanges
window.GR.Sig.All$score <- -10*log10(window.GR.Sig.All$FDR)
window.GR.Sig.Up$score <- -10*log10(window.GR.Sig.Up$FDR)
window.GR.Sig.Down$score <- -10*log10(window.GR.Sig.Down$FDR)
# Add names to the GRanges
names(window.GR.Sig.All) <- paste0("window.all", 1:length(window.GR.Sig.All))
names(window.GR.Sig.Up) <- paste0("window.up", 1:length(window.GR.Sig.Up))
names(window.GR.Sig.Down) <- paste0("window.down", 1:length(window.GR.Sig.Down))
# Export GRanges within RSE objects as BED files
export(window.GR.Sig.All, paste0(exportDir, analysis, "_windows_sigAll.bed"))
export(window.GR.Sig.Up, paste0(exportDir, analysis, "_windows_sigUp.bed"))
export(window.GR.Sig.Down, paste0(exportDir, analysis, "_windows_sigDown.bed"))

# Significant adjacent windows within 50bp of one another are merged (clustered) together into larger regions up to 5000bp
# Merge significantly differential windows based on direction
Merged.Up <- mergeResults(
    winResults.Sig.Up, tab = rowData(winResults.Sig.Up),
    tol = windowSize, merge.args = list(max.width = 5000)
)
Merged.Down <- mergeResults(
    winResults.Sig.Down, tab = rowData(winResults.Sig.Down),
    tol = windowSize, merge.args = list(max.width = 5000)
)
# Create GRanges objects from merged data
Merged.Sig.Up <- Merged.Up$regions
Merged.Sig.Down <- Merged.Down$regions
# Add a score to the GRanges objects
Merged.Sig.Up$score <- -10*log10(Merged.Up$best$FDR)
Merged.Sig.Down$score <- -10*log10(Merged.Down$best$FDR)
# Add names to distinguish them for union
names(Merged.Sig.Up) <- paste0("region.up", 1:length(Merged.Sig.Up))
names(Merged.Sig.Down) <- paste0("region.down", 1:length(Merged.Sig.Down))
# Create a union of directional significant merged regions
Merged.Sig.All <- GenomicRanges::union(Merged.Sig.Up, Merged.Sig.Down, ignore.strand = FALSE)
# Add names to that union
names(Merged.Sig.All) <- paste0("region.all", 1:length(Merged.Sig.All))
# Export merged regions
export(Merged.Sig.All, paste0(exportDir, analysis, "_merged_sigAll.bed"))
export(Merged.Sig.Up, paste0(exportDir, analysis, "_merged_sigUp.bed"))
export(Merged.Sig.Down, paste0(exportDir, analysis, "_merged_sigDown.bed"))

# Save image of workspace
save.image(file = paste0("workspace/", analysis, ".RData"))

# Exit R
q()


#######################
# ### Example of file structure for sample annotation
# ### Named "My_ATAC_Files_Shifted.txt"
### Beginning of file

# Sample	Group	Replicate	BAM_Folder	Extension	BAM_Path
# ATAC_C2C12_Grow_r1	Grow	1	~/My_project_name/bam_and_bai_files/	.star_index_noAnnot.markedDups.ATACshift.bam	~/My_project_name/bam_and_bai_files/ATAC_C2C12_Grow_r1.star_index_noAnnot.markedDups.ATACshift.bam
# ATAC_C2C12_Grow_r2	Grow	2	~/My_project_name/bam_and_bai_files/	.star_index_noAnnot.markedDups.ATACshift.bam	~/My_project_name/bam_and_bai_files/ATAC_C2C12_Grow_r2.star_index_noAnnot.markedDups.ATACshift.bam
# ATAC_C2C12_Grow_r3	Grow	3	~/My_project_name/bam_and_bai_files/	.star_index_noAnnot.markedDups.ATACshift.bam	~/My_project_name/bam_and_bai_files/ATAC_C2C12_Grow_r3.star_index_noAnnot.markedDups.ATACshift.bam
# ATAC_C2C12_Grow_r4	Grow	4	~/My_project_name/bam_and_bai_files/	.star_index_noAnnot.markedDups.ATACshift.bam	~/My_project_name/bam_and_bai_files/ATAC_C2C12_Grow_r4.star_index_noAnnot.markedDups.ATACshift.bam
# ATAC_C2C12_Diff_r1	Diff_T24	1	~/My_project_name/bam_and_bai_files/	.star_index_noAnnot.markedDups.ATACshift.bam	~/My_project_name/bam_and_bai_files/ATAC_C2C12_Diff_r1.star_index_noAnnot.markedDups.ATACshift.bam
# ATAC_C2C12_Diff_r2	Diff_T24	2	~/My_project_name/bam_and_bai_files/	.star_index_noAnnot.markedDups.ATACshift.bam	~/My_project_name/bam_and_bai_files/ATAC_C2C12_Diff_r2.star_index_noAnnot.markedDups.ATACshift.bam
# ATAC_C2C12_Diff_r3	Diff_T24	3	~/My_project_name/bam_and_bai_files/	.star_index_noAnnot.markedDups.ATACshift.bam	~/My_project_name/bam_and_bai_files/ATAC_C2C12_Diff_r3.star_index_noAnnot.markedDups.ATACshift.bam

### End of file
